# Effects of hybridization and gene flow on gene co-expression networks

**DOI:** 10.1101/2024.12.04.626860

**Authors:** Runghen Rogini, Daniel I. Bolnick

**Affiliations:** Department of Ecology and Evolution, University of Connecticut, Storrs, USA

**Keywords:** gene co-expression networks, hybridization, gene flow, network evolution

## Abstract

Gene co-expression networks are a widely used tool for summarizing transcriptomic variation between individuals, and for inferring the transcriptional regulatory pathways that mediate genotype-phenotype relationships. However, these co-expression networks must be interpreted with caution, as they can arise from multiple processes. Here, we show that hybridization and gene flow between populations can greatly modify co-expression networks. Admixture between populations produces correlated expression between genes experiencing linkage disequilibrium. This correlated expression does not reflect functional relationships between genes, but rather depends on migration rates and physical linkage on chromosomes. Given the prevalence of gene flow between divergent populations in nature, these introgression effects likely represent a significant force in network evolution, even in populations where hybridization is historical rather than contemporary. These findings emphasize the critical importance of considering both evolutionary history and genomic architecture when analyzing gene co-expression networks in natural populations.

## 1 Introduction

Understanding the genotype-phenotype relationship remains a fundamental challenge in biology. High-throughput sequencing has revealed that most phenotypic traits arise from complex interactions among multiple genes (Boyle et al., 2017; Mackay et al., 2020). This polygenic architecture manifests through both sequence variation and differential gene expression (Liu et al., 2019; Albert and Kruglyak, 2018). Genome-wide association studies (GWAS) demonstrate that trait variation typically involves many loci acting concurrently (Visscher et al., 2017; Weigel and Nordborg, 2015), and arises from polymorphism in both coding and regulatory sites. An important area of research today concerns the relationship between those many genes. Do they interact epistatically or independently? To what extent is gene expression coordinated across many genes underlying polygenic traits? Molecular, cellular, and developmental biology all reveal extensive networks of gene regulation, in which transcription factors and suppressors alter the expression of other genes, sometimes producing long chains of regulation spanning many genes (Peter and Davidson, 2015; Barabási and Oltvai, 2004). Therefore a key step in understanding genotype-to-phenotype mapping is the description of gene regulatory networks (Wagner, 2011; Payne and Wagner, 2019).

One approach to infer gene regulatory networks, has been the statistical estimation of co-expression between genes (Stuart et al., 2003; van Dam et al., 2018). If a transcription factor *A* causes a target gene *B* to be expressed, then individuals with higher (or lower) expression of *A* will have correspondingly higher (lower) expression of *B* (Langfelder and Horvath, 2008; Zhang and Horvath, 2005). We can measure this association by quantifying how expression of A varies among individuals, how B varies, and the correlation between A and B (Serin et al., 2016). Scaling up to a transcriptome, we can build a correlation matrix between many genes (Butte et al., 1999; Butte and Kohane, 2000). We expect that transcription factors’ expression will be positively correlated with the expression of the genes they up-regulate, and inhibitors’ expression negatively correlated with the expression of their targets (Marbach et al., 2012; Roy et al., 2014). Reversing this logic, biologists often use evidence of correlated gene expression to infer that the genes in question are directly interacting in a regulatory cascade.

Interpreting correlated expression as evidence of gene regulation presents significant challenges. False-negatives can occur when transcription factors show minimal expression variation or exhibit non-linear relationships with target genes (Marbach et al., 2012; De Smet and Marchal, 2010). Conversely, false-positives can arise from various mechanisms unrelated to direct regulation: chromatin accessibility patterns (Dong et al., 2020), shared upstream regulators (Gilad and Mizrahi-Man, 2019), cell-type specific expression variation (Crow et al., 2016), and genetic linkage (Albert and Kruglyak, 2018). Therefore, co-expression modules require careful interpretation as indicators of functional relationships (Langfelder and Horvath, 2008; Serin et al., 2016).

In this paper, we identify a previously overlooked source of co-expression between functionally unrelated genes (Stuart et al., 2003; Langfelder and Horvath, 2008; Serin et al., 2016). In natural populations, gene flow between genetically divergent populations is common (Goulet et al., 2017; Martin et al., 2013; Nosil et al., 2009; Payseur and Rieseberg, 2016). Gene flow can also occur between recently diverged species that can hybridize, enabling introgression of heterospecific haplotypes into another species’ genome. Here, we show that gene flow and hybridization can substantially alter gene-gene co-expression networks. These evolutionary processes can create blocks of highly correlated gene expression that might be identified as modules in a gene co-expression network, but which may have no shared biological function. These blocks exist because hybridization produces linkage disequilibrium (LD) between physically linked genes on a given chromosome. That LD between two or more genes, each with varying expression, produces gene-gene co-expression, that is gradually broken down by recombination across many generations. We used individual-based models to simulate hybridization and gene flow between populations with genetic differences in average gene expression (e.g. with divergent eQTL). Overall, our study shows that gene–gene correlations tended to increase with hybridization, and even in cases of low migration rate. As such, our results suggest that gene co-expression modules may be created by chromosome structure, recombination, and evolutionary history. This provides yet another reason why co-expression modules may not be directly informing us about regulatory networks and gene functional relationships.

## 2 Materials and methods

We used individual-based simulations to model the effect of hybridisation and introgression on gene coexpression networks (Figure 1) under two scenarios: In scenario 1, we simulate a hybrid mapping population, such as might be produced experimentally for Quantitative Trait Locus (QTL) mapping. We simulate an initial F1 hybrid cross between two genetically divergent parental populations, followed by successive generations of sibling matings between hybrids to produce F2, F3, and later hybrid generations (Figure 1 (B)). This simulation would also apply to a hybrid zone between two populations or species. In the second scenario, we model the impact of recurrent immigration into a large resident population (Figure 1 (C)). This mimics the widespread process of gene flow between divergent populations, to evaluate the potential effect of gene flow on gene co-expression networks in natural settings.

**Figure 1.**
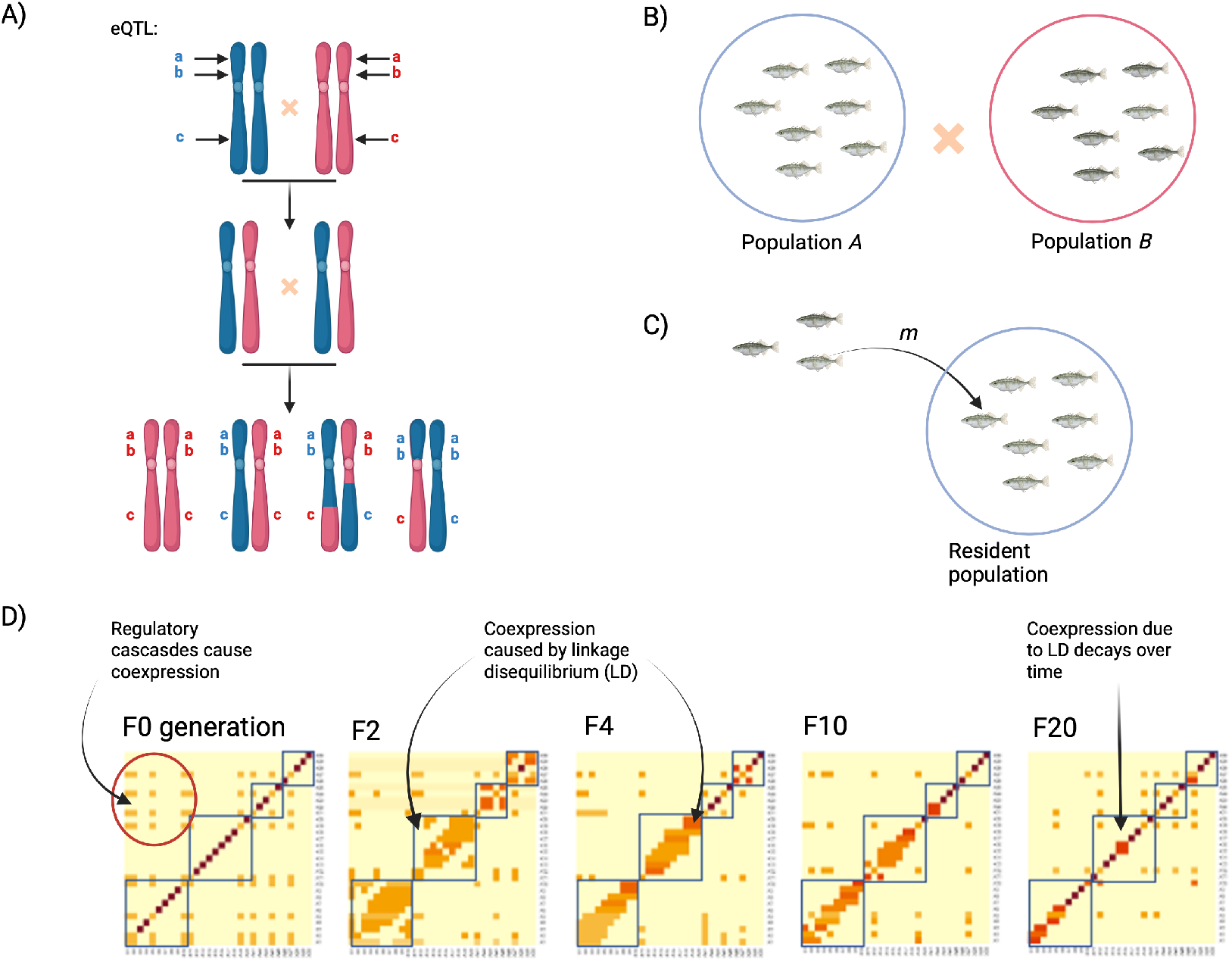
Schema of individual based model used for simulating hybridization and gene flow with recombination events. A) Breeding between two individuals generating F1 and F2 hybrids. Note that *a, b, c* represent cis-acting eQTLs. B) Scenario 1: Hybrid mapping population—breeding occurs between individuals from two divergent parental populations: Population *A* and *B*. C) Scenario 2: Gene flow between two divergent populations. Resident Population *A* receives migrants at a low migration rate *m*. D) A Gene Co-expression Network (GCN) is inferred for the hybrids generated. Genetic modules are estimated to identify co-expressed genes for each GCN inferred.

### 2.1 Scenario 1: Simulating symmetric hybridization (e.g., QTL Mapping Populations)

#### 2.1.1 Parental populations and breeding

We begin with two parental populations characterised by fixed allele frequency differences *(F*_*ST*_ *≈* 1) at numerous loci around the genome. Each individual is a diploid with *N*_*chr*_ chromosomes, and the *i*^*th*^ chromosome contains *N*_*i*_ loci with fixed differences (e.g. five chromosomes with 10, 10, 5, 5, and 10 loci respectively). Population *A* is fixed for alleles with value 0 at these loci (e.g., genotypes 0/0, and Population *B* is fixed for alleles with value 1 at these loci (genotypes 1/1). We then assume that there is a hybrid mating between one individual from population *A*, and one individual from population *B*. Each parent contributes one haploid genome to generate an F1 hybrid that is a heterozygote (genotype 0/1) at all loci. The F1 population is assembled with *N* individual offspring from this mating (here, *N* = 500 except where noted). To generate the F2 hybrid generation, we randomly selected two F1 hybrid individuals (with replacement) to pair up and produce an F2 offspring. Each individual F1 parent generates a haploid gamete with a single recombination event per chromosome per generation. The chromosomes undergo independent assortment to produce a gamete that has a mix of 0 and 1 alleles in linkage blocks. Recombination is modelled by using a uniform random number to select the position of a crossing over event within a given chromosome, choosing one chromatid’s contents to the left and the other chromatid contents to the right of the cross-over. Recombined chromatids from each parent are used to create a new diploid individual, an F2 hybrid, and this is repeated to again reach a population size of *N* F2 offspring. The process is repeated by sampling (with replacement) pairs of F2 individuals to be parents of F3 offspring, and so on for 20 generations of hybrid crossing.

#### 2.1.2 Simulating the transcriptome

In any given generation, each individual organism’s genotype is converted into a transcriptome as follows. Each locus has a genetic value *G*_*j*_ of 0 (genotype 0/0), 1 (genotype 0/1) or 2 (genotype 1/1) that may impact its own expression level (e.g. cis-regulatory variation). Each locus *i* is randomly assigned a mean expression rate *α*_*i*_ ∼ 𝒩 (*μ*_*α*_, *σ*_*α*_), and a cis-eQTL effect size *β*_*i*_ ∼ 𝒩 (0, *σ*_*β*_). The expected expression rate of the gene *i* in individual *j* (*λ*_*ij*_) is a logit function of the individual’s genotype, *λ*_*ij*_ = *logit*(*α*_*i*_ + *β*_*i*_ *∗ G*_*j*_). The observed expression level of gene *i* in individual *j, E*_*ij*_ is then drawn from a Poisson distribution with expectation *λ*_*ij*_. This generates locus-specific mean, variance, and genotype-dependence of expression levels to simulate varying effect size cis-eQTL throughout for all loci.

Next we generate an underlying gene co-expression network architecture representing true regulatory cause-effect relationships. We use a Barabási-Albert preferential attachment algorithm (Barabási and Albert, 1999) (power scale of 1.0, implemented in the igraph package (Csárdi et al., 2024)) to create a network. To ensure that only a subset of genes are involved in a causal regulatory network, we multiply each element of this simulated adjacency network by a Bernoulli random variable with a user-defined network density *D*. This ensures that most pairs of genes are uncorrelated in any causal sense. We add weights by multiplying all non-zero elements of the adjacency matrix connecting gene *i* to gene *k* by an effect size unique to each pair of genes *γ*_*ik*_ *∼ N* (0, *σ*_*γ*_). Then, we take the uncorrelated transcriptome (described above, and for each pair of genes *ik* that are connected in the network, we generate a new expression level for a focal gene *i*, that depends on the randomly generated expression of that gene *i*, but is also modified by expression of a second gene, *k*. Hence, expression of gene *i* in individual *j* is:

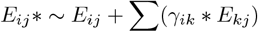

to produce positively or correlated gene pairs (depending on the sign of *γ*_*ik*_).

To incorporate genotype-dependent networks, we add the possibility that one master gene modifies the structure of the rest of the network. The two parental populations can be given independently generated co-expression networks (e.g. each ancestral population has a different matrix of weights *γ*_*ij*_ to allow for population differences in gene regulatory controls). This difference is controlled by one randomly chosen locus. In F1 and later generations, each individual’s genotype at the master locus determines which network is applied to generate a simulated transcriptome for that individual. For heterozygotes at the master locus, the network is the average of the edge strengths (including 0’s for absent edges) of the two parental types. Simulations were also conducted using a single regulatory architecture for the two parental populations, but giving them different mean expression levels.

#### 2.1.3 Analysis

We ran the stochastic individual-based simulations described above to generate populations of parents, F1 hybrids, F2 hybrids, F3 hybrids, through the F20 generation. As this is meant to be a simulation to demonstrate overall trends, we used a simplified genome of five chromosomes with 5-10 loci each. Transcriptome parameters were chosen to generate fairly realistic variation in overall gene expression levels, co-expression, and eQTL effect sizes (*μ*_*α*_ = 5, *σ*_*α*_ = 2, *σ*_*β*_ = 2, *σ*_*γ*_ = 2, network density *D* = 0.2, and network parameters *σ*_*γ*_ = 2.5 and density *D* = 0.1 in the second population). These effect sizes correspond closely to empirical eQTL effects estimated from a recent F2 hybrid mapping population of threespine stickleback (Fuess et al., 2021; Weber et al., 2022; Hu et al., 2024). For each generation we obtained the transcriptomes of *N* = 500 individuals and used a threshold correlation coefficient of | *r* |*>* 0.2 to define the presence of edges in the coexpression network. We plotted gene-gene co-expression networks for each generation and tracked changes in the degree distribution, modularity, and transitivity. We repeated this with the parental populations having identical underlying regulatory networks, and with separate regulatory networks.

### 2.2 Scenario 2: Simulating gene flow between divergent populations

#### 2.2.1 Resident and immigrant parental populations

We begin with two parental populations characterized by fixed allele frequency differences at numerous loci around the genome. Each individual is a diploid with *N*_*chr*_ chromosomes, and the *i*^*th*^ chromosome contains *N*_*i*_ loci with fixed differences (five chromosomes with 10, 10, 5, 5, and 10 loci respectively). The focal population *A*—the resident population—is fixed for alleles with value 0 at these loci (genotypes 0/0), and Population *B*—the migrant population—is fixed for alleles with value 1 at these loci (genotypes 1/1). We model directional gene flow from population *B* into population *A*, which maintains a constant size of 500 individuals except where noted.

Migration follows a Poisson process, where the number of migrant individuals (*N*_*m*_) arriving in each generation is drawn from a Poisson distribution with expectation *m*. Under this model, the expected fraction of immigrants in population *A* each generation is 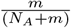. We examine a detailed gradient of migration rates by varying *m* on a logarithmic scale, sampling 1001 evenly spaced points from exp(*−*10) to 1 (specifically, *m* = exp(*x*) where *x* ranges from -10 to 0 in increments of 0.01). This fine-grained sampling allowed us to detect subtle transitions in population genetic and network properties across different immigration regimes. The model represents a continent-island scenario with unidirectional gene flow, focusing on the dynamics of the island population.

#### 2.2.2 Breeding

At each generation, mating occurs at random amongst the individuals in population *A*. That is, in the first generation mating can either occur between two individuals from the resident population, or between a resident and an immigrant individual, or between two immigrants (if there are enough of them). We do not simulate sexual selection or natural selection against immigrants here. In the very first generation, if a resident mates with an immigrant, they create an F1 hybrid who is heterozygous at all loci. These hybrids can then reproduce with each other, or with homozygous residents or immigrants (mating remains random with respect to genotype or phenotype). In subsequent generations, the resident population becomes a hybrid mixture of different generations of backcrosses, with perpetual input of new immigrants from population *B*.

Each generation proceeds as follows: first, Poisson-distributed immigrants arrive from population *B*; then, random mating pairs are drawn (with replacement) to produce offspring until the target population size of 500 is reached. Each parent contributes a haploid gamete formed through recombination, with one recombination event per chromosome per generation (similar to Scenario 1). The recombination process uses a uniform random number to select a breakpoint within each chromosome, with the resulting gamete containing one chromatid’s contents to the left of the breakpoint and the other chromatid’s contents to the right. Recombined chromatids from each parent create a new individual, and this is repeated to again reach a population size of *N* .

#### 2.2.3 Genetic architecture and gene expression

Each individual’s genotype affects its gene expression through both cis- and trans-regulatory variation. The transcriptome is modeled with parameters chosen to generate realistic variation in expression levels, coexpression patterns, and eQTL effect sizes. Each locus *i* is assigned a mean expression rate *α*_*i*_ *∼ N* (*μ*_*α*_, *σ*_*α*_), where *μ*_*α*_ = 5 and *σ*_*α*_ = 2, and a cis-eQTL effect size *β*_*i*_ *∼ N* (0, *σ*_*β*_), where *σ*_*β*_ = 3.

The gene regulatory networks differ between the parental populations, with population *A* having network density *D* = 0.02 and effect size *σ*_*γ*_ = 2, and population *B* having density *D* = 0.03 and effect size *σ*_*γ*_ = 1.5. Similar to Scenario 1, one randomly chosen locus serves as a master regulator that controls which network architecture is expressed. In the resident population *A* and immigrant population *B*, individuals homozygous at this locus express their respective population’s network architecture. In hybrid individuals, the genotype at this master regulatory locus determines which network is expressed: homozygotes express their respective parental network architecture, while heterozygotes express a network with edge weights averaged between the two parental networks. This allowed us to track how network architecture evolves under gene flow, as the regulatory control itself can introgress between populations along with other loci.

#### 2.2.4 Analysis

We ran numerical simulations for 100 generations at each migration rate, recording both genetic and transcriptomic outcomes. For genetic analysis, we track allele frequencies at all loci across generations (refer to the Supplementary Information for further details). For transcriptome analysis, we obtain expression profiles for all 500 individuals after 100 generations of introgression. Gene–gene co-expression networks are constructed using a threshold correlation coefficient of | *r* |*>* 0.2 to define edges. We analyzed network properties including mean degree distribution, modularity, and transitivity to characterize how gene regulatory architecture changes with varying levels of gene flow. Multiple independent replicates are run for each parameter combination to assess stochastic variation in outcomes.

The simulation framework is implemented in R (R Core Team, 2024), using parallel processing to efficiently run multiple replicates. Network visualization and analysis are performed using the igraph package (Csárdi et al., 2024), with custom functions for calculating network metrics and generating comparative visualizations across different migration rates.

## 3 Results and Discussion

In order to identify previously overlooked source of co-expression between functionally unrelated genes in natural populations, here we used a individual-based simulation approach to hybridization (Scenario 1) and gene flow (Scenario 2) between two divergent populations. Overall, both of our simulations show that both hybridization and gene flow can substantially alter gene-gene interactions. More importantly, these evolutionary processes can create blocks of highly correlated gene expression that might be identified as modules in a gene co-expression network, but which may have no shared biological function. Rather, the modules can be an artefact of linkage between functionally unrelated but co-inherited cis-eQTL.

### 3.1 Hybridization restructures gene co-expression networks across multiple generations

Our simulations confirmed that gene co-expression networks are altered by the process of hybridization. Even first generation hybrids tended to have modestly different networks than either parental population (Figure 2, Figure S2). This difference between parental and F1 networks is a consequence of starting with genetically divergent parental populations with distinct networks, and assuming additive effects in the F1s. As a result F1s will have more edges overall, the union of the set of non-zero edges in each parental population. However, edge strength will tend to be weaker, being the average of the parental populations. This result is sensitive to assumptions about dominant (or additive) effects of loci that alter gene regulatory network structure. Alternatively, if we define the parental populations as having identical regulatory network edges and weights, then the F1 hybrid network is the same as the parents (Figure S1). Mean expression of any one gene will still be different (intermediate) in the hybrids than in the parents, because they are heterozygous for cis-regulatory effects. But, these differences in mean do not alter whether or not there is co-expression between genes. Thus, the first generation of hybridization may produce a distinct and more connected network, if parental networks are different to begin with (refer to SI for other examples of F1 genotypes simulations Figure S2). The specifics of this F1 hybrid network depend on assumptions about genetic dominance coefficients of the edge strengths.

**Figure 2.**
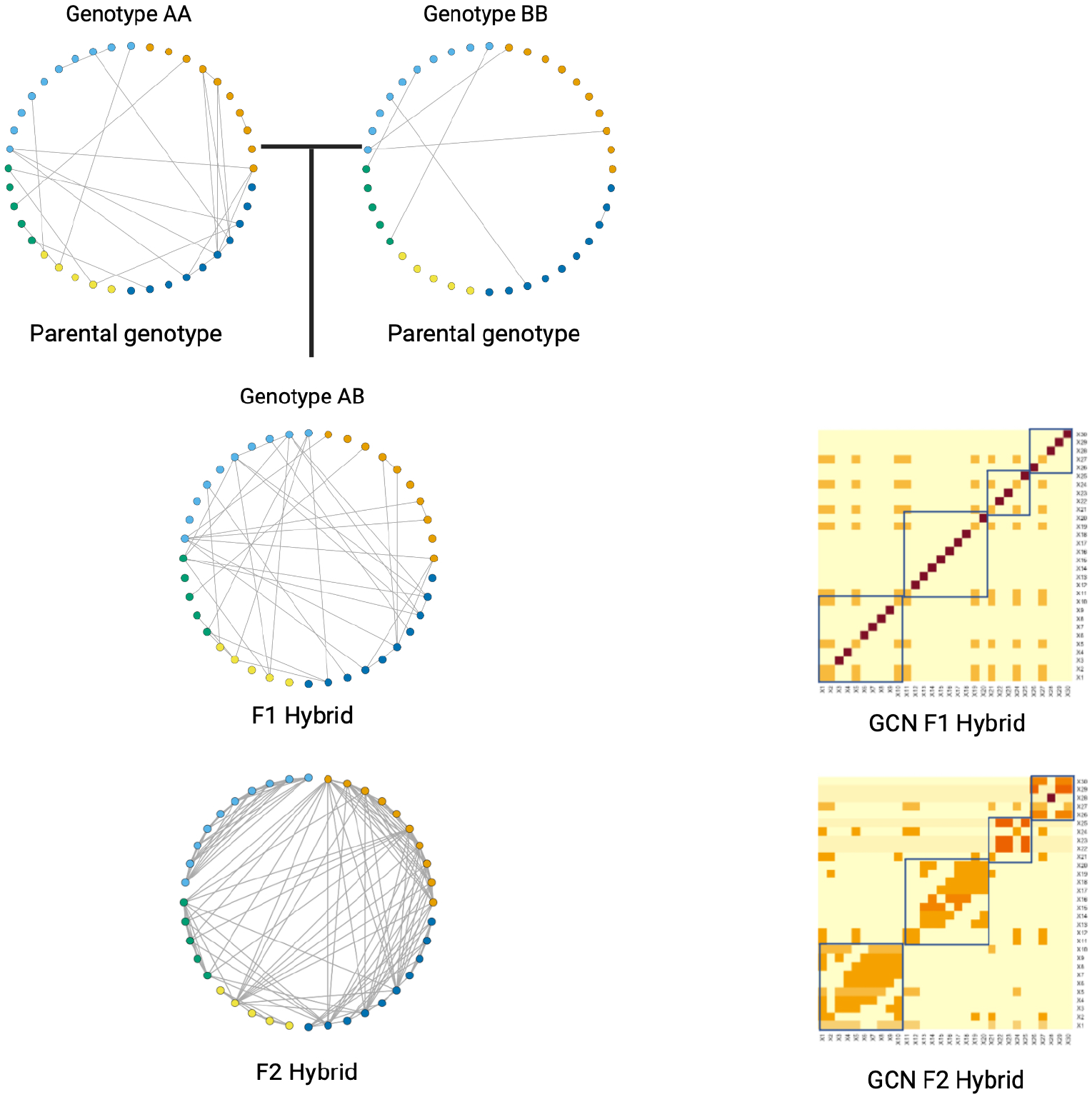
A gene-gene interaction network inferred from QTL Mapping simulation (Scenario 1). The parental genotype *AA* and *BB* generates F1 hybrids i.e. genotype *AB*. The F1 hybrids subsequently generate F2 hybrids (example co-expression network from simulations are presented here). In the gene-gene interaction network of both the F1 and F2 hybrids, there is an increase of gene-gene interactions with hybridization. As observed by the Gene Co-expression Networks (GCN), genetic modules are formed in the F2 hybrid as a result of co-inheritance of cis-eQTL from adjacent loci. Note that transcripts are color coded by chromosome in the circular plots.

Recombination during F1-F1 intercrosses led to a substantial increase in network density (e.g. mean degree) in the F2 hybrid generation (Figure 2). This is because the low rate of recombination causes correlated inheritance of cis-eQTL from physically adjacent loci along a given chromosome. Individuals inheriting a 0 allele at gene *i*, are more likely to also inherit a 0 allele at gene (*i* + 1). The result is large genetic blocks (haplotypes) spanning sections of chromosomes. When converted into a simulated gene expression network, these polymorphic haplotype blocks create modules of co-expressed genes that correspond to whole chromosomes (e.g., Figure 2). Even the correlation between genes on different chromosomes become denser. This happens because upstream regulatory genes in the network are co-inherited with other genes on the same linkage block, so all these linked genes become correlated with the downstream targets of their neighboring regulatory gene. For instance, gene X27 (Figure 2) on chromosome 4 is correlated with multiple loci on chromosome 1, because there exists a single upstream regulatory gene on chromosome 1 that targets X27, but other chromosome 1 genes are inherited with and so correlated with this regulatory element.

Hybrid gene regulatory networks exhibit a distinct three-phase evolutionary trajectory, driven by the interplay between hybridization and recombination. Initially, hybrids experience substantial network disruption, marked by a dramatic spike in connectivity during the F2 generation, showing an almost 5-fold increase in mean degree compared to parental networks. This initial disruption phase is followed by a gradual reorganization period, where recombination in subsequent generations begins breaking up physical linkage blocks responsible for correlated gene expression.

The network then enters a stabilization phase, characterized by enhanced organization compared to parental genotypes. Although mean degree decreases by approximately half by the F3-F4 generation and continues declining due to progressive recombination breaking linkage blocks, the hybrid networks maintain elevated connectivity (approximately two-fold higher) even after 20 generations. Importantly, these networks also exhibit higher modularity and transitivity values compared to both parental and F1 populations. Hybridization thus produces sustained changes in correlated gene expression, which might incorrectly be used to infer a gene regulatory architecture. This effect decays very gradually because linkage disequilibrium takes a very long time to decay.

This persistent elevation in network metrics has two significant implications. First, it demonstrates that genetic linkage plays a fundamental role in shaping gene co-expression network evolution, with effects lasting many generations beyond the initial hybridization event. Second, it suggests that observed patterns of co-expressed genes may, in some cases, result from physical linkage rather than functional regulatory relationships. These findings reveal how hybridization and subsequent recombination events can create longlasting changes in regulatory network architecture, potentially contributing to hybrid fitness through modified gene regulatory organization. Furthermore, it is interesting to speculate that these sustained changes in correlated expression might create opportunities for changes in regulatory circuits, and may be a target of selection. In the present paper we have not considered the effects of natural selection acting on these linkage blocks, treating gene expression (and correlations) as effectively neutral, but this would be a valuable next direction.

### 3.2 Introgression and LD shape gene co-expression networks

Gene flow substantially influences co-expression network architecture through a complex relationship with migration rate. The effects manifest in three key network metrics: connectivity (mean degree), local clustering (transitivity), and modularity. At moderate migration rates (*>*1e-02 migrant individuals per generation on average), networks show increased connectivity and local clustering, while highest rates (approaching 1 migrant per generation) enhance modularity, demonstrating how gene flow can restructure transcriptional networks.

The impact of recurrent immigration reveals a non-linear relationship with network density. At very low migration rates, limited introgression occurs as immigrants are typically lost through genetic drift, leaving networks largely unchanged from resident populations. However, successful introgression of genomic blocks in some cases increases network degree above both donor and resident population levels. As migration rates increase, stochastic variance decreases, showing more consistent effects across replicates.

Notably, even after 100 generations, populations receiving moderate immigration maintain higher network connectivity than both resident and immigrant populations. This persistence occurs through LD between adjacent loci, creating apparent trans-acting effects between chromosomes without functional basis. For example, increased co-expression between genes on different chromosomes can result from LD between regulatory elements and their target genes.

At the highest migration rates, a slight decline in mean degree occurs as immigrant genotypes increasingly replace the resident population, trending toward a single population network structure, though retaining elevated connectivity due to persistent LD. These findings demonstrate how hybridization, immigration, and genetic linkage collectively shape gene co-expression network evolution, with effects that can persist long after the initial genetic exchange.

## 4 Concluding remarks

Our simulations demonstrate that hybridization and introgression fundamentally alter gene co-expression networks through complex interactions between genetic linkage and regulatory architecture. The effects persist across multiple generations, with network changes manifesting through both cis- and trans-regulatory relationships. Importantly, our results reveal that observed co-expression patterns in hybrid populations may often reflect physical linkage rather than functional regulatory relationships, particularly in regions of strong linkage disequilibrium. This has crucial implications for interpreting gene co-expression networks in hybrid systems, as distinguishing between correlation due to genetic linkage versus genuine regulatory interactions becomes challenging. These findings emphasize the importance of considering both evolutionary history and genomic architecture when studying gene co-expression networks, particularly in populations experiencing hybridization or ongoing gene flow. Gene flow is common among many divergent populations (and even species) in nature. Therefore, the introgression effects described here have the potential to be a major force driving evolution of gene co-expression networks. The fact that migration and hybridization can have a sustained effect, means that this concern affects populations even if introgression and gene flow are periodic, and historical, rather than contemporary. Studies of gene co-expression networks must therefore keep in mind the potential for gene flow and linkage to be an important process.

## Data Availability Statement

All data in this manuscript were simulated, and all corresponding codes for the simulations are publicly available on Zenodo (Runghen and Bolnick, 2024) and GitHub (https://github.com/rogini98/gcn_hybridization.git).

## Authors’ contributions

DIB designed this study, both DIB and RR contributed to the code of both simulations and analysis. DIB and RR contributed to the writing, editing of the current manuscript.

## Acknowledgements

RR and DIB acknowledge funding from the National Science Foundation (FAIN-2133740).

## Supplementary Information

### Impact of hybridization on gene co-expression networks

**Figure S1:**
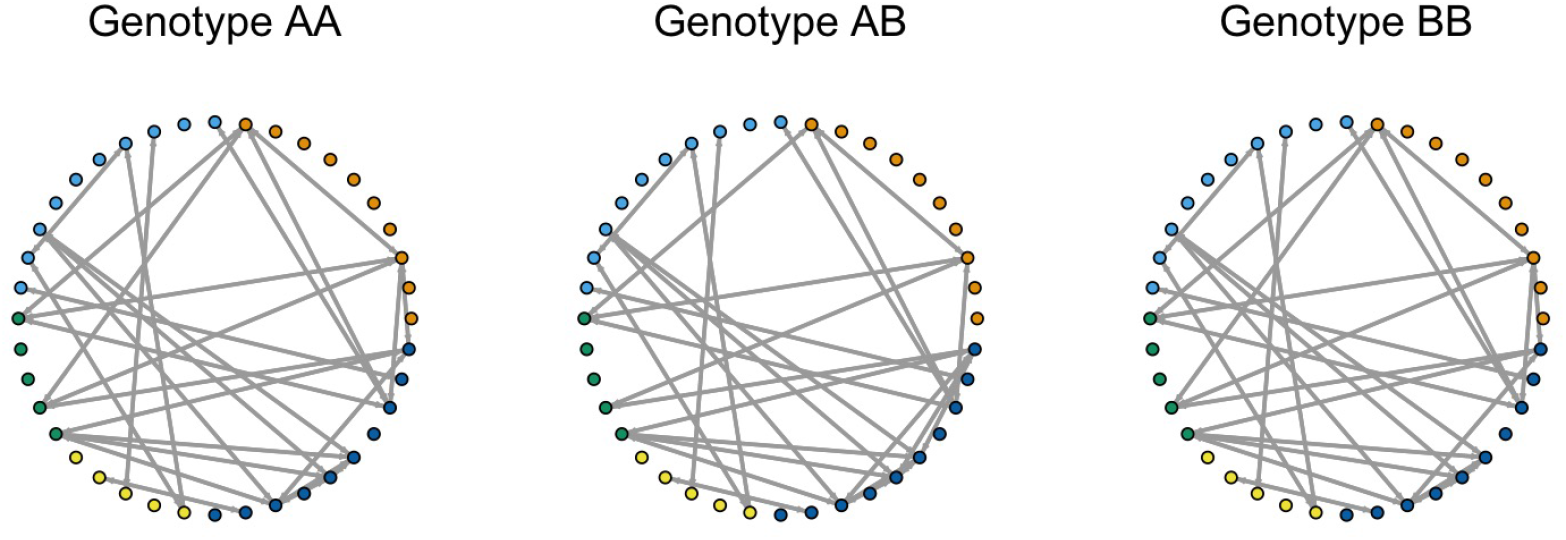
Conservation of network structures across parental and hybrid genotypes. An identical network topology is observed across all three genotypes. This demonstrates a complete conservation of the network architecture when parents share identical regulatory elements and expression patterns.

### Gene-gene interaction network of F1 genotype from individual based simulation of Scenario 1

**Figure S2:**
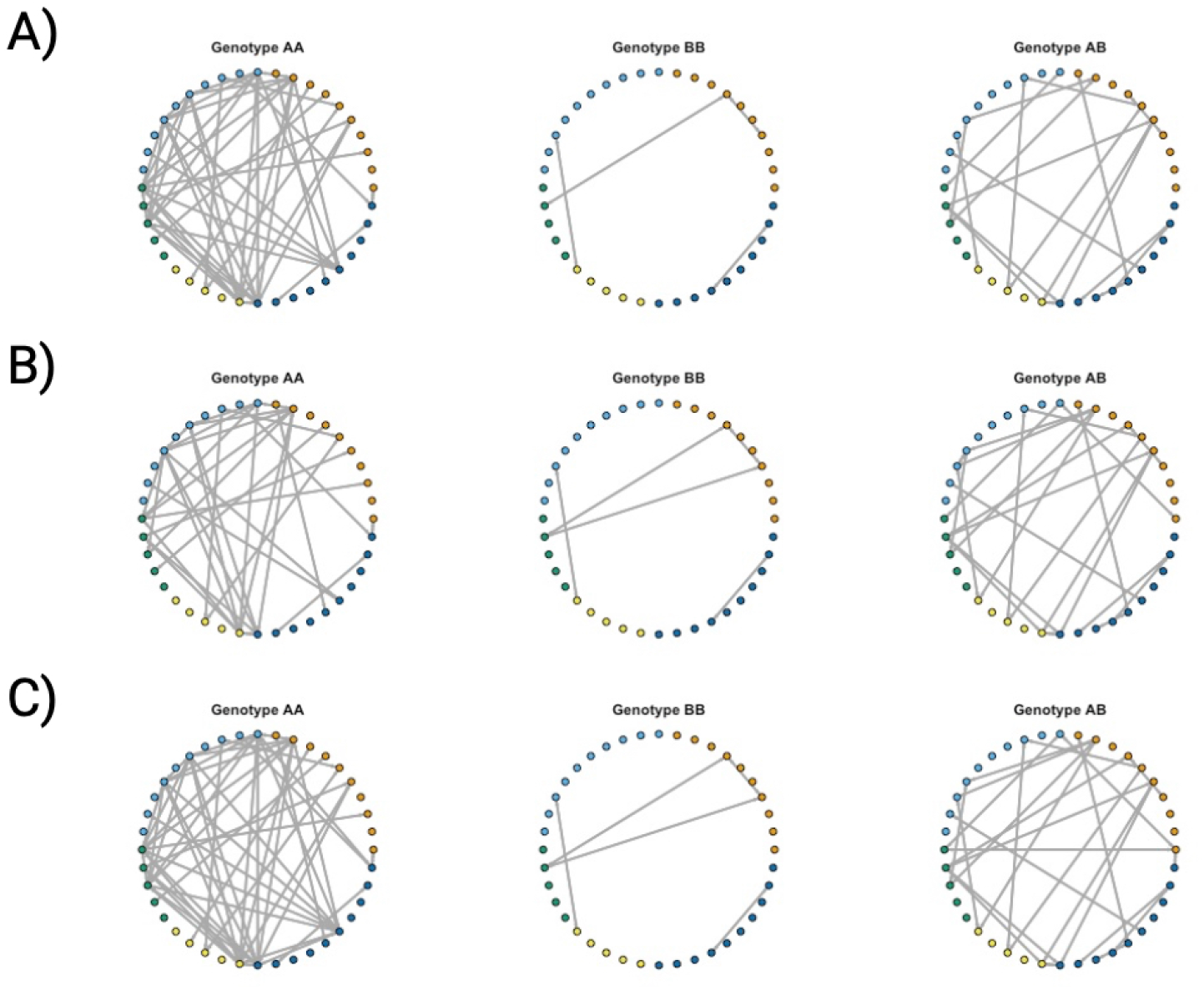
Gene-gene interaction network inferred from QTL Mapping simulation (Scenario 1). The parental genotype AA and BB generates F1 hybrid i.e. genotype AB.

### Comparing the network metrics relationships in hybrid Gene Co-expression networks

**Figure S3:**
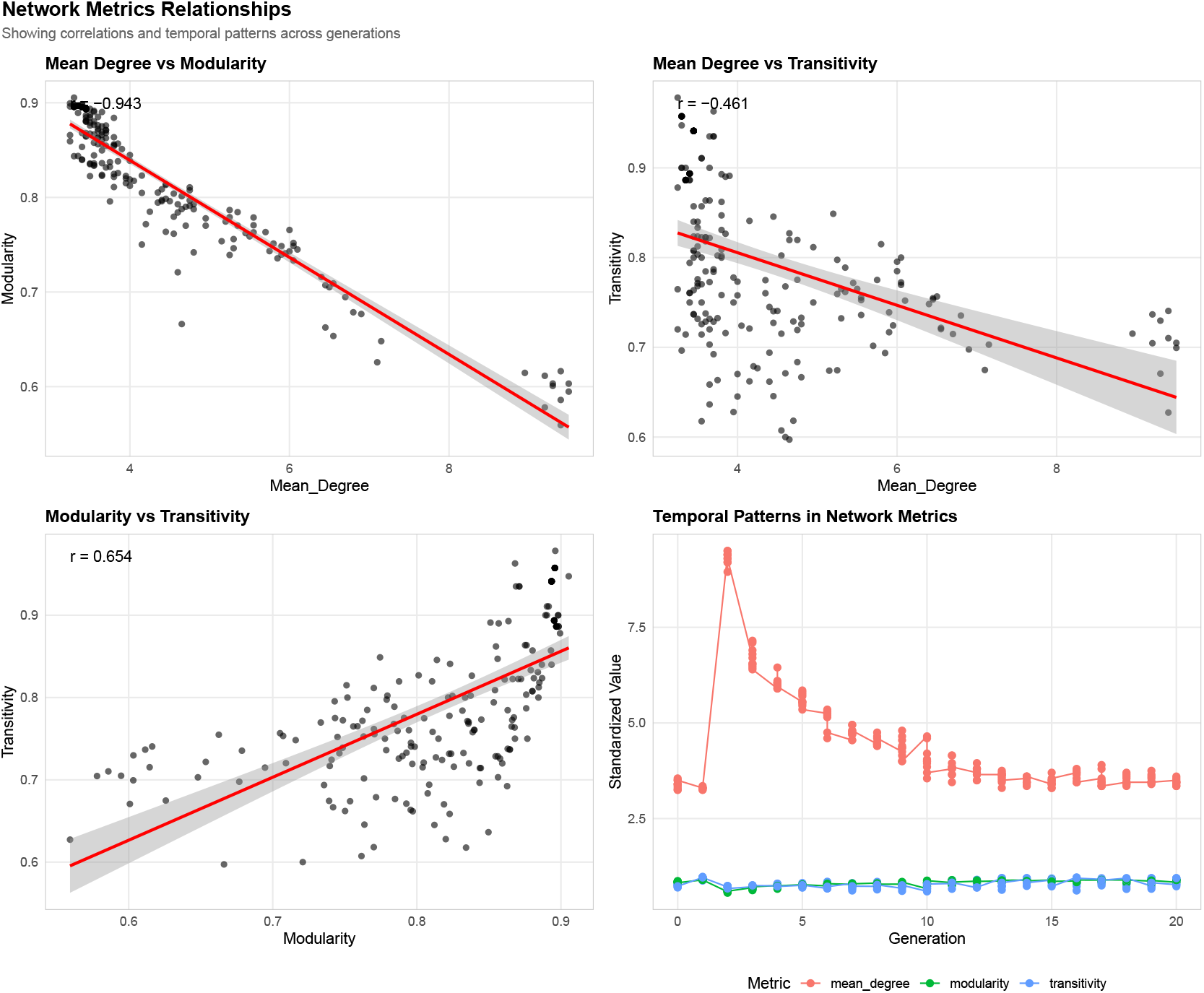
Network metric relationships across hybrid generations. Pairwise correlations between network metrics (mean degree, modularity, transitivity) and their temporal evolution over 20 generations. A strong negative correlation between mean degree and modularity is observed amongst hybrid generations. In contrast, a positive modularity-transitivity relationship is observed across the hybrid generations. Overall, while looking at the temporal patterns of the network metrics, an initial network disruption followed by stabilization is observed. Note that the grey bands indicate 95 % confidence intervals.

### Tracking allele frequency evolution under variable migration rates

#### Migration and breeding design

We simulated introgression scenarios with variable migration rates (*m ∈* 0, 0.05, 0.5, 1) between two populations fixed for alternative alleles. The resident population (*A*) was initialized with allele value 0 at all loci (genotypes 0/0), while the migrant population (*B*) was fixed for allele value 1 (genotypes 1/1). Each generation, the number of migrants (*N*_*m*_) was drawn from a Poisson distribution with expectation *m*. For each migration rate, we conducted *n* = 10 replicate simulations to account for stochastic variation in the introgression process.

#### Population structure and breeding

The population size was maintained at *N* = 500 individuals per generation. In each generation, after migration occurred, random mating was implemented by sampling parent pairs with replacement from the combined resident and migrant pool. Each parent contributed gametes generated through independent assortment of chromosomes, with one recombination event per chromosome. This process was repeated for 100 generations to allow sufficient time for allele frequencies to reach equilibrium under the different migration regimes.

#### Tracking and analysis

For each replicate simulation at each migration rate, we tracked: 1) generation-by-generation allele frequencies at each locus, 2) the final population structure after 100 generations, 3) complete transcriptome profiles incorporating both direct genetic effects and network-mediated expression patterns. The expected expression rate of gene *i* in individual *j* (*λ*_*ij*_ ) was modeled as:

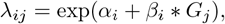

where *α*_*i*_ ∼ 𝒩 (5, 2) represents the baseline expression rate, *β*_*i*_ ∼ 𝒩 (0, 5) represents the effect size, and *G*_*j*_ is the genetic value (0, 1, or 2). For each migration rate scenario, we calculated: the mean allele frequencies across replicates, the standard deviation of allele frequencies, 95% confidence intervals derived from quantiles of the replicate distribution (Figure S4)and network metrics (including mean degree, transitivity, and modularity - Figure 3).

**Figure 3.**
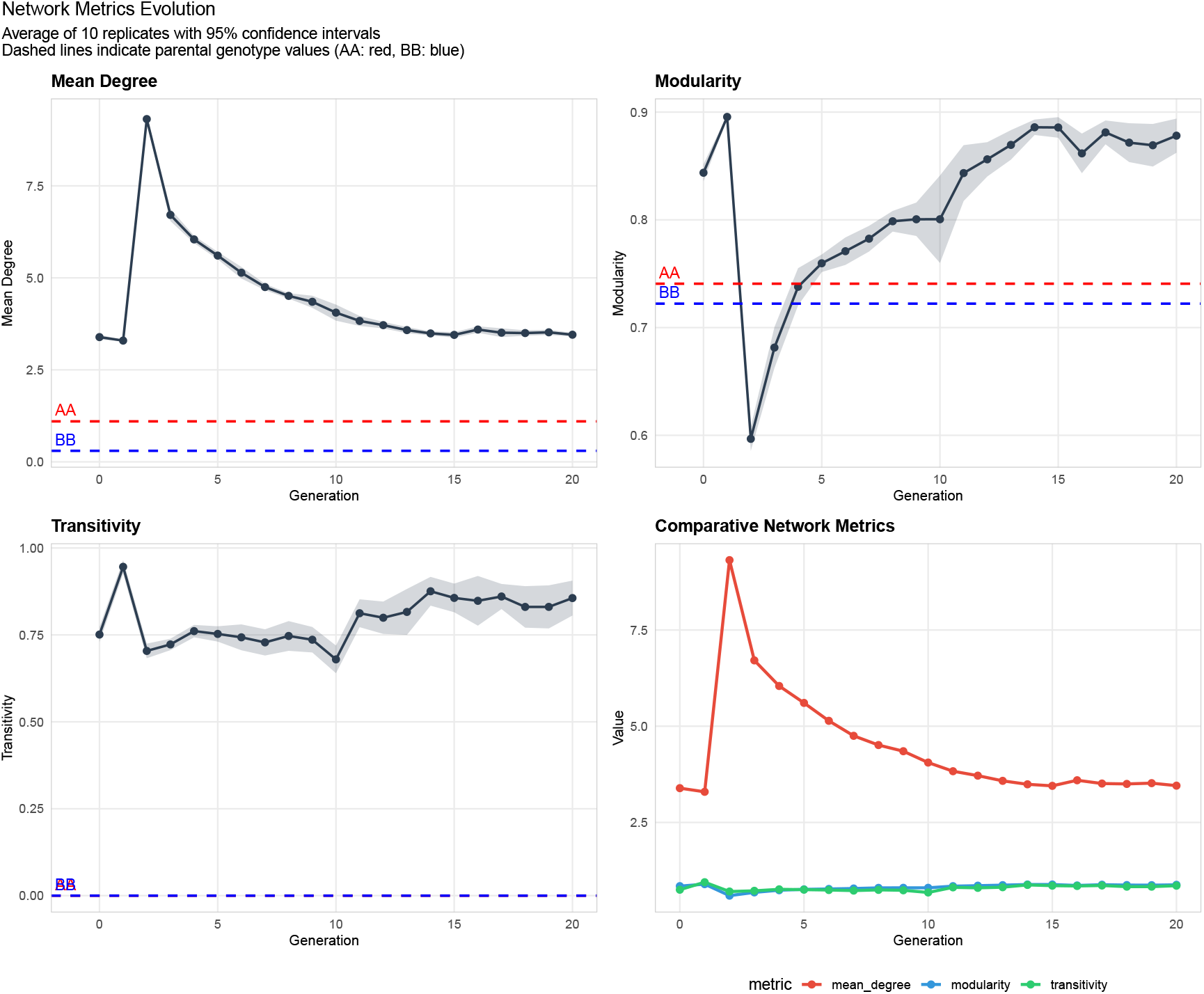
Temporal evolution of hybrid network architecture. Network metric evolution averaged across 10 replicates with 95 % confidence intervals (shaded grey areas). Dashed lines represent parental genotype values (*AA*: red, *BB*: blue). Mean Degree shows initial network disruption followed by stabilization above parental values. Modularity demonstrates early decrease (F3 – F5), but recovers and exceeds parental levels, indicating new genetic module organization. Transitivity indicates enhanced local clustering of genes compared to parental genotypes. Network metric comparison—bottom right panel—highlights the trade-off between connectivity (mean degree) and network organization (modularity and transitivity) during hybrid network evolution. This temporal change highlights how hybrid regulatory networks initially experience disruption but ultimately might achieve a potentially new architectural state which is distinct from both parental genotypes.

**Figure 4.**
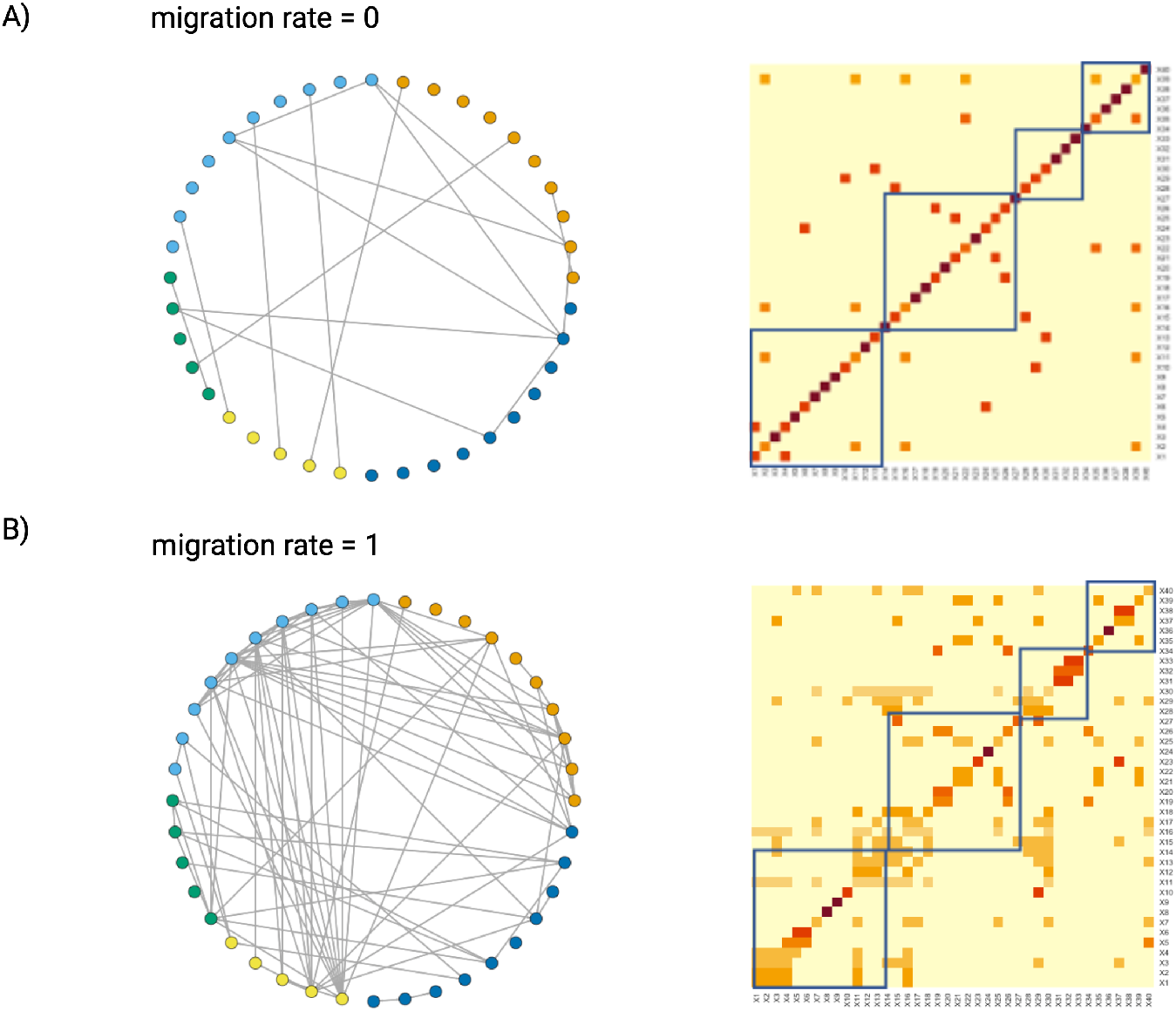
Gene Co-expression Networks generated from simulation run with rare migration over 100 generations (averaged from 10 replicates). A) GCN inferred with no migration shows that the gene-gene association in the resident population is low, reflecting causal gene regulation effects; B) GCN inferred from a resident population receiving one migrant per year (on average) shows an increasing number of gene-gene interactions and correlation caused by an increase in correlated inheritance of cis-eQTL from adjacent loci.

**Figure 5.**
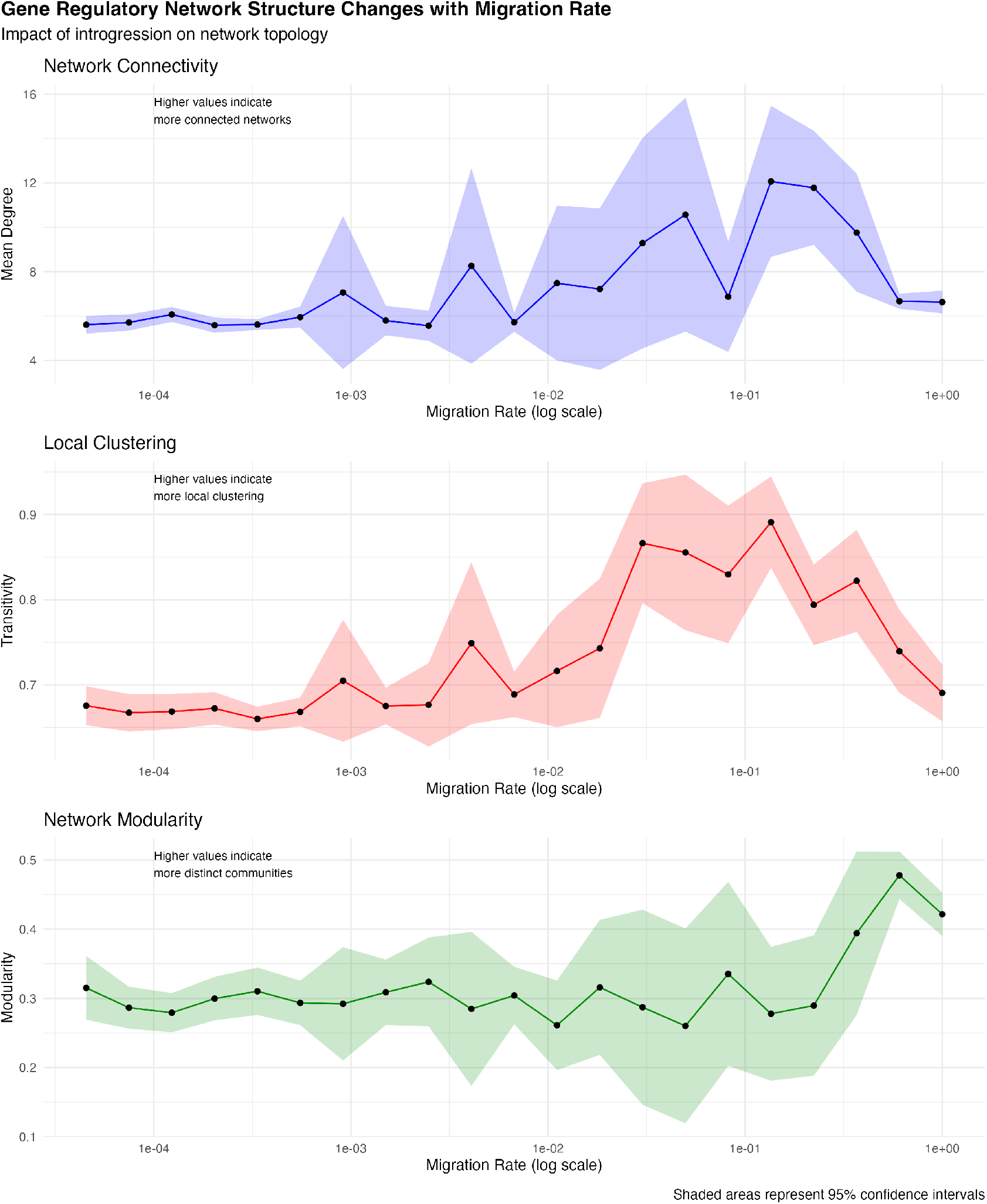
Impact of introgression on the network topology of the Gene Regulatory network inferred at F100 generation hybrids at different migration rates (Model from Scenario 2). We tracked the mean degree, transitivity and modularity of the gene co-expression. High values of mean degree indicate more connected networks, high values of transitivity indicate more local clustering and high values of modularity indicate more distinct communities. Note that the shaded area represents the 95 % confidence intervals.

**Figure S4:**
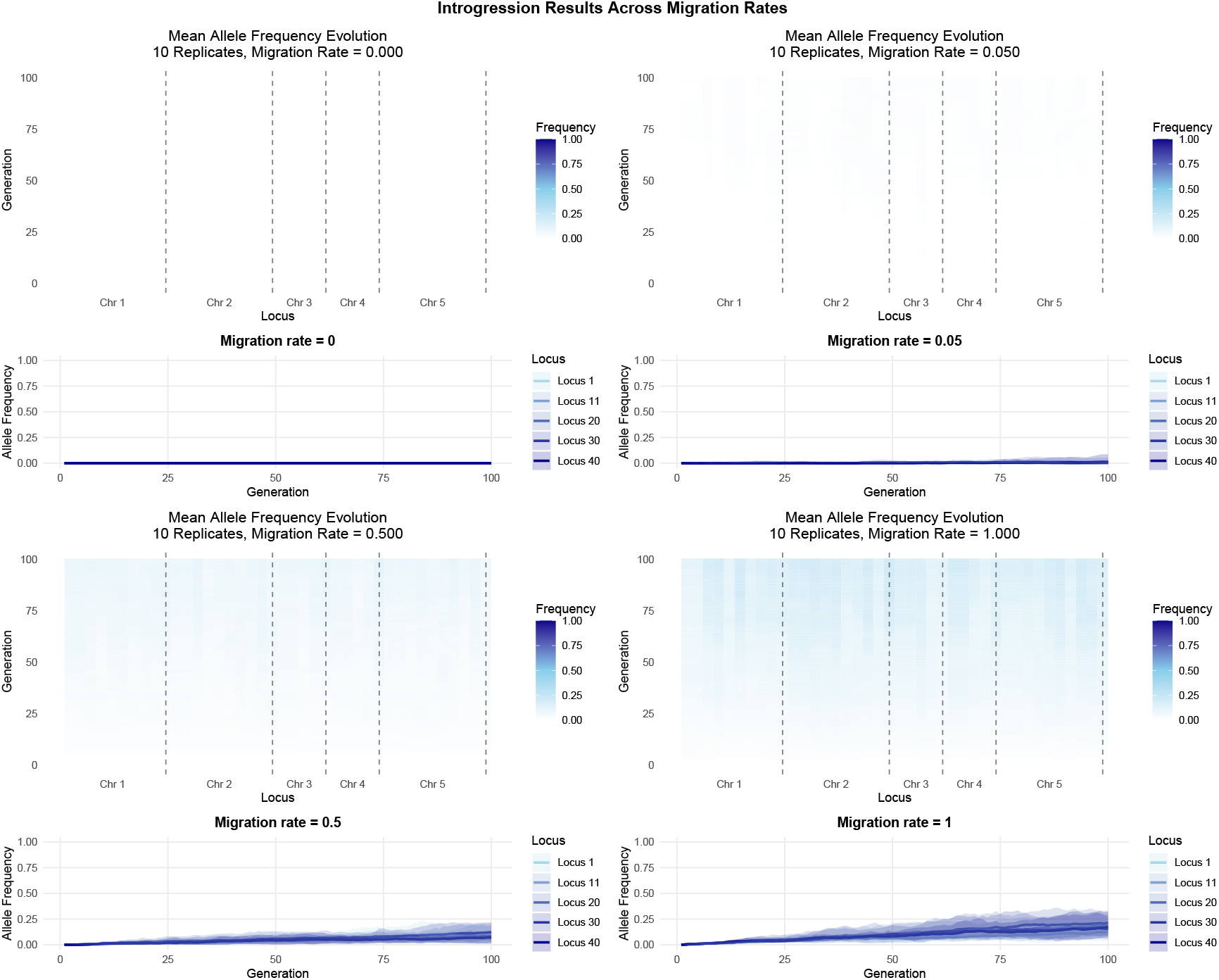
Migration rate effects on genomic introgression patterns. Four-panel comparison showing allele frequency dynamics under different migration rates (0 – 1) over 100 generations (averaged over 10 replicates). Heatmaps (top) show mean allele frequency evolution across chromosomal regions (Chr 1-4) over 100 generations. Trajectory plots (bottom) show allele frequency trajectories with 95 % confidence intervals for five tracked loci. The color gradient (white to dark blue) indicates allele frequency from 0 to 1. Higher migration rates (0.5 and 1) show increased allelic introgression across chromosomal regions. Increasing migration rates show corresponding increases in genomic admixture and allelic introgression. Note that the vertical dashed lines in heatmaps denote chromosomal boundaries.

### Null Model: Control for Network Evolution

#### Parental populations and breeding

We implemented a simplified genetic architecture to serve as a null model control. Each individual is a diploid with *N*_*chr*_ = 50 chromosomes, with each chromosome containing exactly one locus (*N*_*i*_ = 1 for all *i*) to eliminate physical linkage effects. Population *A* is fixed for alleles with value 0 at these loci (genotypes 0/0), and Population *B* is fixed for alleles with value 1 (genotypes 1/1). Following initial hybridization, we generated an F1 population of *N* = 500 heterozygous individuals (genotype 0/1 at all loci). For subsequent generations (F2 through F20), we randomly selected two individuals (with replacement) to pair up and produce offspring. Each parent contributed one haploid gamete, with independent assortment of chromosomes ensuring random mixing of alleles. This breeding process was repeated to maintain a constant population size of *N* = 500 individuals per generation.

#### Simulating the transcriptome

For each individual organism, the genotype was converted into a transcriptome as follows. Each locus *i* was assigned a mean expression rate *α*_*i*_ ∼ 𝒩 (5, 2) and a cis-eQTL effect s ize *β* _*i*_ ∼ 𝒩 (0, 5 ). T he expected expression rate of gene *i* in individual *j* (*λ*_*ij*_ ) was calculated as:

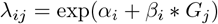

where *G*_*j*_ represents the genetic value of 0 (genotype 0/0), 1 (genotype 0/1), or 2 (genotype 1/1). To implement network effects, we generated three independent network matrices with density *D* = 0 .1. Each non-zero element in these matrices was assigned an effect size *γ* _*ik*_ ∼ 𝒩 (0, 2 ). A single randomly chosen locus served as the network eQTL, determining which network matrix was active for each individual based on their genotype at this locus. For heterozygotes at the network eQTL, effects were averaged between matrices. The final expression value for each gene was constrained to b e non-negative:

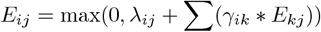

#### Analysis

We tracked changes in network structure across 20 generations of hybrid crossing. For each generation, we obtained transcriptomes of *N* = 500 individuals and used a threshold correlation coefficient of | *r* |*>* 0.2 to define the presence of edges in the co-expression network. We calculated three key network metrics: 1) Mean degree (average connectivity between genes), 2) transitivity (local clustering coefficient) and modularity (community structure assessed via fast greedy clustering). To verify the absence of linkage disequilibrium, we calculated pairwise correlations between loci at generations 0, 5, 10, and 20. All simulations were conducted with a fixed random seed (123) to ensure reproducibility. Parameters were chosen to match the scale of variation observed in previous hybrid mapping studies (*σ*_*α*_ = 2, *σ*_*β*_ = 5, *σ*_*γ*_ = 2, network density *D* = 0.1).

**Figure S5:**
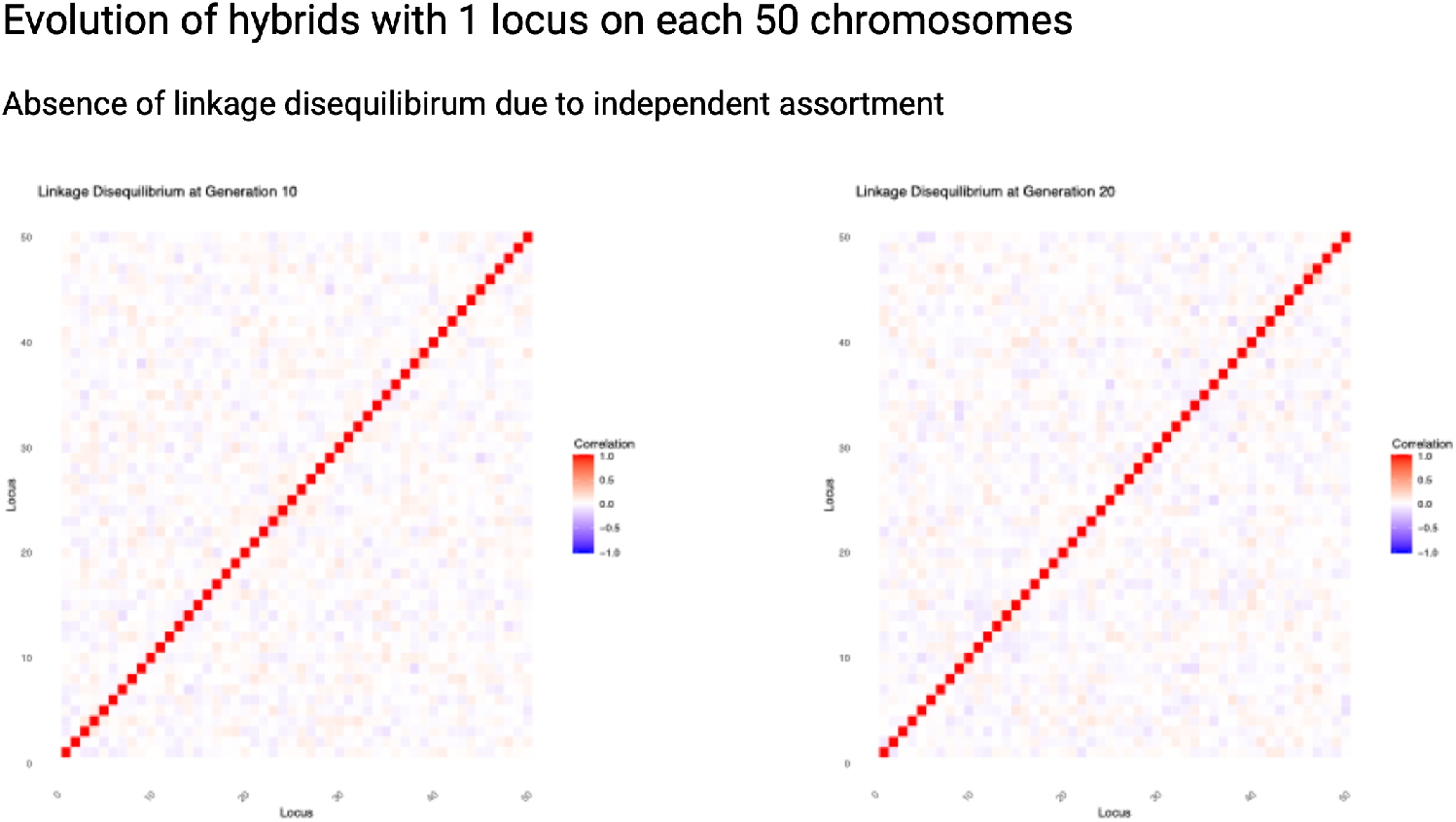
Linkage Disequilibrium (LD) patterns in null model. Absence of LD in hybrid populations simulated with 50 chromosomes, each containing one locus, tracked over 20 generations. The two heatmaps compare LD patterns at generation 10 (left) and generation 20 (right). The correlation values range from *−*1 (blue) to 1 (red). The strong diagonal red line (correlation = 1) represents self-correlation of loci, while the predominantly white/pale background (correlation *≈* 0) across off-diagonal elements indicates absence of LD between loci on different chromosomes. This pattern confirms independent assortment of alleles across the 50 chromosomes, which persists across the hybrid generations.

## Notes

https://github.com/rogini98/gcn_hybridization.git

### Competing Interest Statement

The authors have declared no competing interest.

